# Analysis of MRC-supported early career researcher success in obtaining follow-on research funding

**DOI:** 10.1101/2020.02.19.949263

**Authors:** Ian Viney, Buddhini Samarasinghe, Kevin Dolby

## Abstract

This study examined the progress of MRC-funded researchers, with a focus on their success in securing subsequent research income. We report here results from compiling details of the research income of researchers that completed an MRC early career award between 2011 and 2015. Depending on the funding scheme 50-59 per cent of researchers spent on average more than 100,000 GBP per year from new grants secured within three years of finishing their MRC award. In each scheme around 30 per cent of researchers secured less than 50,000 GBP per year research funding in the three years following their MRC award. We apply the same approach to an analysis of progression for the MRC cadre of intramurally-supported Programme Leader Track researchers, and a sample of MRC research grant holders. We also supplement our results with a small number of interviews with researchers that held MRC New Investigator Research Grants. We suggest that these data confirm reasonably good progression rates, with MRC early career schemes demonstrably effective in supporting research careers. They confirm there is a bottleneck with some researchers taking several years to establish their research career, and that expansion of the research base will need balanced new investment across schemes and stages and cannot be driven through growth in early career schemes alone.

## Introduction

### MRC support for early career researchers

The Medical Research Council (MRC), now part of UK Research and Innovation, is the largest UK public funder of discovery biomedical research. The MRC supports research and training across the entire spectrum of medical sciences, for researchers at every career stage, in universities and hospitals, as well as in its own units, centres and institutes in the UK, and in units in Africa^1^.

The MRC is interested in understanding more about how researchers (those supported via dedicated schemes for early career researchers in particular) progress to establish a substantive research programme. This transition to independence is a critical career step and may be made in a variety of ways. Early career researchers may secure funding from the MRC through several different routes:

- Research grant funding won in competition with later career stage researchers via applications considered by the MRC’s research boards or via directed calls for proposals
- Securing a core-funded programme leader-track position in an MRC Institute or Unit
- Career Development Awards (CDAs), these are fellowships for non-clinical (and clinically inactive) researchers
- Clinician Scientist Fellowships (CSFs) for clinical researchers
- New Investigator Research Grants (NIRGs), these are awarded via the MRC’s research boards, and are prioritised for consideration during funding to support researchers securing their first research grant.

The MRC is interested in striking the right balance between specialised schemes to support the transition to independence and general funding for research awarded by its research boards and via directed calls. The MRC is concerned by reports that early career researchers find it difficult to build a sustained academic research programme.

### Evidence from previous analyses

The US National Institutes of Health (NIH) often refers to the career progression from early career award to the next substantive competitively won grant, as “K2R”. This is where a researcher transitions from a “K” type NIH award (targeted to supporting the transition to independence), to R01-type project grants (open to the whole research population).

A large NIH evaluation of K award holder outcomes^2^ examined 792 mentored early career K grant holders, and their subsequent success in securing NIH funding between 2000 and 2005. The proportion that graduated from a K-type grant to obtaining any other NIH grant ranged between 57% (for K23 grant holders) to 50% (for K01 grant holders). The RO1 grants are between 1 and 5 years in duration (average 4.5), no distinction was made in the scale of follow-on funding achieved by these researchers. This study exclusively used NIH funding data and noted that there was a need to look across all biomedical funders when analysing funding portfolios.

An older study^3^ of US National Cancer Institute (NCI) supported early career researchers obtained details of subsequent funding awards from a wider set of funders (public and charitable supporters of cancer research in the US). This study analysed 1,200 NCI K-grant holders and found that 56% secured some subsequent research grant funding. However, the scale of subsequent support was not examined.

Other studies have examined the influence of gender^4,5,6,7^, indicating that there were disparities in success between male and female researchers that required further investigation. A survey of UK early career researchers that became principal investigators between 2012 and 2018 found that most (80 per cent of men, 77 per cent of women) secured some grant funding within five years of starting their early career award^8^.

We believe that our study is the first study to compile complete funding details for a set of researchers to allow scale and timing of subsequent support to be investigated.

## Methods

We compiled details of the research income secured by all researchers who, between 2011 and 2015, completed New Investigator Research Grants (NIRG), Clinician Scientist Fellowships (CSF), and non-clinical Career Development Awards (CDA). These special schemes are targeted at providing researchers with their first independent research grant. Similar numbers of researchers at an early career stage are supported by the MRC outside of special schemes, through research grants or directed funding calls. As a comparator we looked at a small sample of these grant holders with similar characteristics (RG). We also analysed the progress of similar researchers in MRC Institutes and Units compiling data on the research income for all programme leader track (PLT) researchers that left their positions between 2011 and 2015). Details about the current position for a small number of early career researchers could not be located and so were excluded from the analysis, this included six researchers that moved from PLT positions soon after taking them up.

**Table 1.** Characteristics of MRC early career researcher awards included in the analysis.

|  | **CDA** | **CSF** | **NIRG** | **RG** | **PLT** |
| --- | --- | --- | --- | --- | --- |
| **Male/Female ratio\*** | 30/16 | 31/10 | 68/41 | 37/9 | 36/12 |
| **% clinically qualified** | 0% | 100% | 8% | 22% | 21% |
| **Average annual award value (£000s)** | 250 | 220 | 140 | 180 | 520 |
| **Median award duration (months)** | 59 | 47 | 36 | 35 | 72** |
| **Median age (years) of researcher at time of completing MRC award** | 39 | 42 | 39 | 41 | 42 |
| **Number included** | 46 | 41 | 109 | 46 | 48 |
| **Number excluded** | 3 | 1 | 7 | 0 | 6 |

\* It should be noted that the analysis was of awards finishing between 2011 and 2015, reflecting applications made 3-5 years prior to this (so awards starting at the earliest 2006 and latest 2012). The gender balance across all these funding routes has changed positively in recent years, meaning that future cohorts studied in the same way will have a greater proportion of female researchers (for example successful applications to the CDA/CSF/NIRG schemes combined in 2017/18 were from 23 male:22 female researchers).
\*\*PLT positions are normally held for six years before a decision is taken about progression to a programme leader post, however start dates were not readily available for this cadre of staff, meaning it was not possible to calculate a median duration for PLT support.

The average annual award values vary significantly between the different funding routes, but some of this can be explained by the fact that CDA, CSF, PLT positions, and (in the period studied here) NIRGs include the full salary costs of the principal investigator, RGs are awards made to Universities at 80 per cent full economic costs, whereas PLT awards are fully funded allocations to MRC units/institutes.

Extensive desk research was undertaken to assemble information about research income. The main sources of research income data we consulted were:

- Researchfish® reports. The MRC requires all its current and recently completed grant holders to provide feedback on the output of MRC funded research via the online Researchfish® system^9^. This feedback includes the option of reporting details of further funding. Over 90 per cent of researchers that held NIRG, CSF or CDAs had submitted responses using Researchfish®. Responses added in the further funding section should answer the question “has your MRC award led to obtaining further funding from the MRC or other sources”, and around 75 per cent of respondents had provided some details in this section. The Researchfish record also indicates if the researcher is still at their original institution and these details are updated annually in discussion with the employing institution.
- Research council grant details. The UKRI Gateway to Research^10^ provided details of research council awards other than the early career award where the researcher is principal or co-investigator.
- Europe PubMed Central (EPMC) grant finder tool. EPMC funders provide grant details to populate the grant finder tool^11^ and this dataset was searched for matches to researcher names and any additional data added.
- University/research organisation websites. Increasingly researcher home pages are populated by University “common research information system” data, and some Universities publish the funding secured by their researchers. Web searches were made for each researcher, and in most cases a relevant home page identified. This was useful to fill some gaps or discover that researchers had moved institution. Universities often publish press releases for major new awards allowing details of award holders to be validated.
- Funder websites. Some funders list awards made by year (e.g. The Leverhulme Trust^12^) which could be consulted for award details. Others provide databases that can be searched online (e.g. US NIH RePORT^13^, Canadian Research Information System (for the Canadian Institutes of Health Research)^14^)
- EU Community Research and Development Information Service (CORDIS)^15^ was consulted for details of European Commission funded awards. The entire CORDIS dataset was downloaded and searched for matches to the names of researchers who had held MRC early career funding.
- Where specific gaps in grant details were identified, but the funder was known, requests were made to the funding organisation to provide data.

We sought to capture the full details of each award (start and end dates, title, funder, and value), in each case we also noted (if the details were available) whether the researcher was the first named/principal investigator or a co-investigator. In the case of networks and consortia funding (e.g. most European Commission funding agreements) a note was made of the funding allocated to the researcher’s organisation in the UK, and the total consortia funding overall.

Where possible data was extracted from an authoritative source (e.g. funder database). In cases where partial data indicated the association of an MRC researcher with an award (e.g. a researcher University webpage listing funding sources with as little detail as “Leverhulme Trust”) other sources were sought to validate this and add enough data for analysis.

From the details of grant and other research income an average annualised research income for the three years following the early career award was calculated.

A small number of semi-structured telephone interviews were conducted with former NIRG holders to validate the research income data that we had compiled and to explore the main barriers to securing follow on funding. The interviewees were asked whether they could highlight factors that assisted or hindered securing research funding and the interview explored whether these factors were part of the researcher’s personal circumstances, were specific to their project or their institution’s environment. Interviewees were asked for their views on how the MRC could better support early career researchers.

The average annual research incomes of different groups of researchers were compared using the Mann-Whitney U test16 and the difference was reported as be significant with a p value higher than 0.05.

Box and whisker charts were produced using the Microsoft Excel template provided by Vertex42^17^.

## Results

### The majority of early career researchers remained in academic research for three years following MRC funding

Most of the 196 ex-CDA/CSF/NIRG researchers in the sample remained in academic roles in the UK for the three years after the MRC award. Twenty moved to research roles in other countries, and two moved to industry research and development roles. Of the 48 PLT researchers, 33 (69 per cent) progressed to MRC programme leader positions in MRC Units/Institutes. The remaining 15 secured group leader/Director positions in academic groups in the UK and overseas. All the researchers in the sample of research grant holders remained in academia.

### The majority of MRC-funded early career researchers secured substantial follow-on research funding

We summarised the proportion of early career researchers securing different levels of research income. We used four ranges (£0-£50k, £50k-£100k, £100k-£400k, and £400k+) for the estimated average annual expenditure from follow-on funding and plotted the number of ECRs in each case achieving this.

CDA, CSF, and NIRG holders all showed similar rates of progression; 50-59 per cent secured funding above £100,000 per year within three years of finishing their career award. These were similar results to the outcomes found in analogous NIH schemes.

In each scheme around a third of researchers (in total 62 out of the 196) secured less than £50,000 per year research funding in the three years following their MRC award. Investigations to identify possible explanations for this found that 40 per cent of the researchers with the lowest category of research income (25 of the 62) had moved institution (which may explain a delay in securing external grant funding), and that 9 of the 25 re-locating researchers had moved to research positions overseas (where complete funding details were also more difficult to verify). There were a few cases from the earlier cohorts in which researchers were successful in gaining grant funding but took longer than three years to do so.

A large proportion of MRC fellows (between 17-20 per cent of CSF and CDA award holders) went on to secure high levels of funding (i.e. above £400,000 per year). Fellows secured a higher average post-award income than NIRGs, although the difference was not found to be significant (comparing the research income of CDA and NIRG holders *p*=0.2). Differences may be due to the longer tenure of fellowships, which allow these researchers an additional 1-2 years to secure a programme of follow-on funding.

The PLT researchers and researchers from the sample that had secured MRC research grants, had a different profile of funding to the ex-CDA/CSF/NIRG holders (p<.0002). For both groups a much higher proportion (between 89 and 94 per cent) of researchers secured more than £100,000 per year research income.

### In the first three years, the largest proportion of research income for ex-MRC funded early career researchers is secured from the MRC

Over 3,000 records of research income were compiled relating to the 290 researchers included in this analysis. Estimated total expenditure from these awards in the three years immediately following completion of each early career award was £367 million. Almost 500 unique sources of funding were identified including UK and international public sector funders, academic institutions, hospital trusts, companies, learned societies, and charities. For approximately two thirds of these awards and roughly half of the funding secured (£180 million) the ex-early career researcher was named as principal investigator.

The MRC remained the largest single source of funding for the population of early career researchers overall (estimated total three-year expenditure of £157 million, 43 per cent of total), and the largest single source of funding for ex-CSF/NIRG/PLT and the sample of research grant holders. For ex-CDA holders the MRC was the second largest funding source (providing a total of £12 million research income for the three years examined, just behind the £13 million secured from Wellcome).

Chart identifies income secured as principal investigator (PI), co-investigator (Co-I), or not determined (N/D).

Ex-CDA/CSF/NIRG holders secured a similar proportion of grants as PI (range 36 – 42 per cent by value and 57 to 62 per cent by number), a higher proportion of ex-PLT researchers secured funding as PI (71 per cent by value, 80 per cent by number), and a lower proportion of ex-RG holders secured funding as PI (35 per cent by value and 43 per cent by value).

### No significant differences found between the research income secured by male and female early career researchers

The progression of male and female researchers was compared across the cohorts of early career researchers. Progression measures for women who held NIRGs were a little lower than for men within three years of finishing the MRC award, with lower overall levels of funding, although the difference was not found to be statistically significant (*p*=0.3). Conversely a small opposite effect was seen for CDAs, where men had a slightly lower median income, although on some other parameters (e.g. numbers achieving very high levels of income) there was no difference. There were no material differences by gender among Clinician Scientist Fellows.

The median income secured by research grant and PLT researchers was two to three times that of the CDA/CSF/NIRG holders. No significant difference was identified between the income of male and female ex-PLT and research grant holders.

In the box and whisker chart the ends of the whisker are set at 1.5 times the interquartile range (IQR) above the third quartile (Q3) and below the first quartile (Q1). If the minimum and/or maximum values are outside these ranges they are plotted as outliers. The number of data points, full number of outliers boundaries between the quartiles are shown in the table below the chart.

### No significant differences found between the research income secured by early career researchers hosted by different research organisations

We investigated whether the employing research organisation for the researcher had any impact on the subsequent success of the early career researcher. We decided to aggregate research organisations into three groups, based on MRC’s approach to strategic relationships with research organisations. Group 1 universities include an MRC strategic investment (i.e. unit or centre) and receive more than or equal to 2 per cent of MRC expenditure on campus. Group 2 universities include an MRC strategic investment and receive between 0.5 and 2 per cent MRC expenditure on campus. The 20 group 1 and 2 organisations receive annual visits from the MRC to discuss research strategy, success rates and research policy issues. The third group includes all other research organisations, including independent research organisations such as public sector and charity funded research establishments.

No significant differences were found between the three groups of institutions, all employed researchers that obtained funding to roughly similar extents, based on the average additional funding gained in the three years immediately following completion of the NIRG/CDA/CSF award. Researchers were no more or less likely to move institution during or immediately after their early career award, if they started their CDA/CSF/NIRG at a Group 1, Group 2 or “other” institution. The data is summarised in the box and whisker plot in Figure 5.

**Figure 1.**
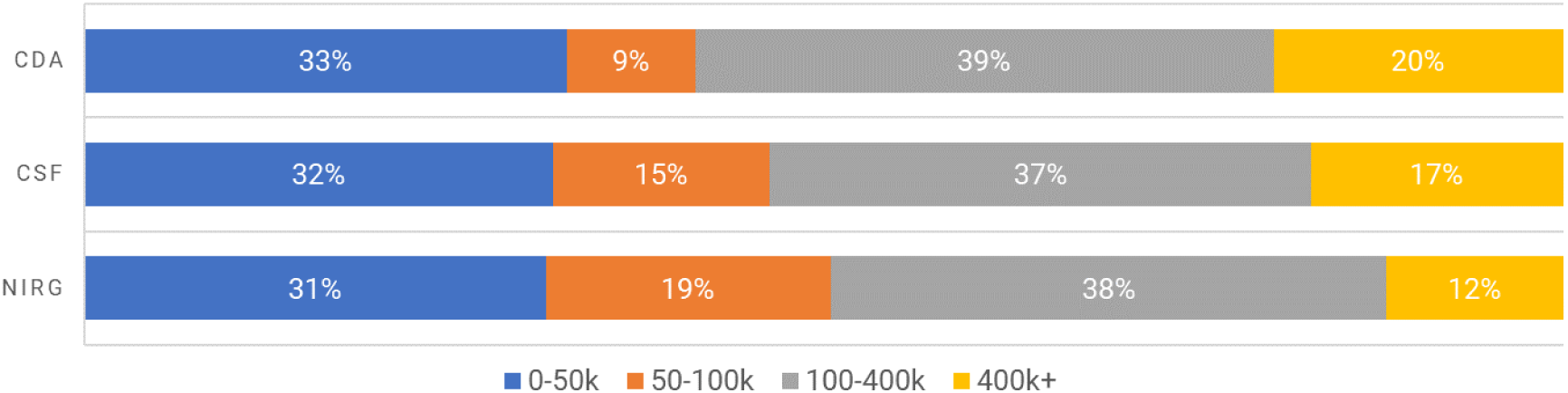
Summary of proportions (%) of researchers achieving different levels of follow-on funding in the <u>three years following</u> their NIRG/CDA/CSF award

**Figure 2.**
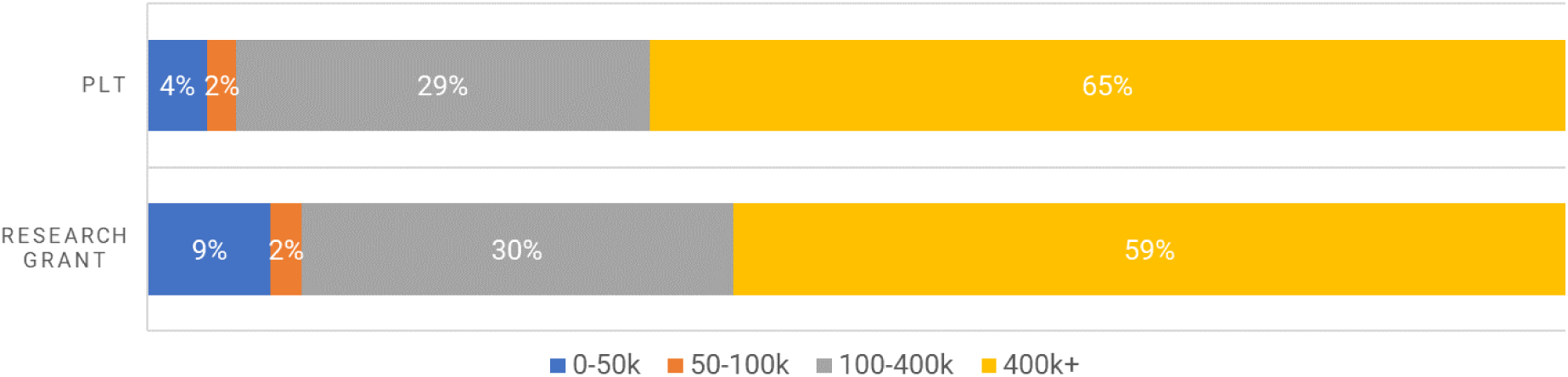
Summary of proportions (%) of researchers achieving different levels of follow-on funding in the <u>three years following</u> completion of their MRC PLT/research grant

**Figure 3.**
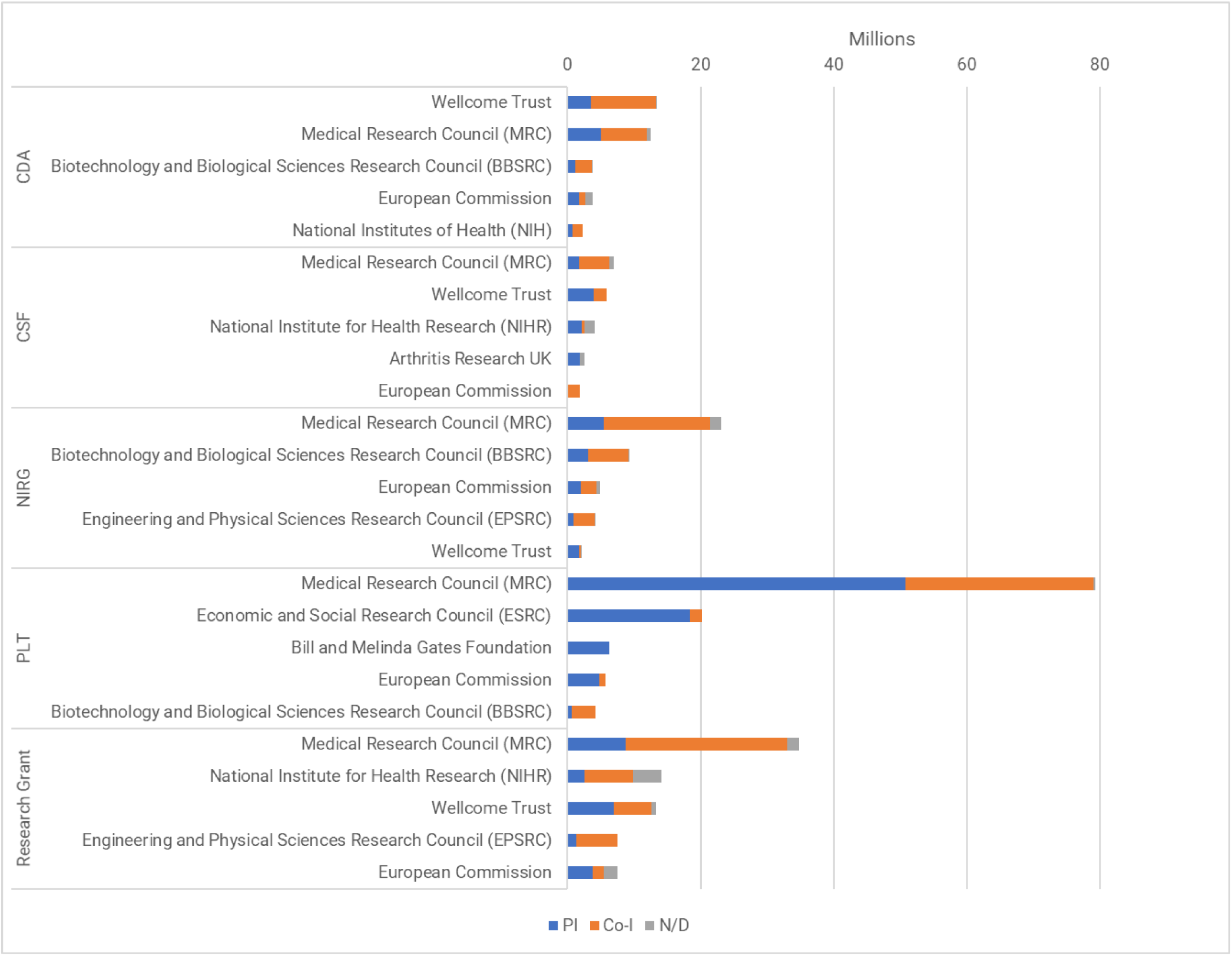
Total research income secured in the three years following completion of CDA/CSF/NIRG/PLT or research grant by top five funding sources

**Figure 4.**
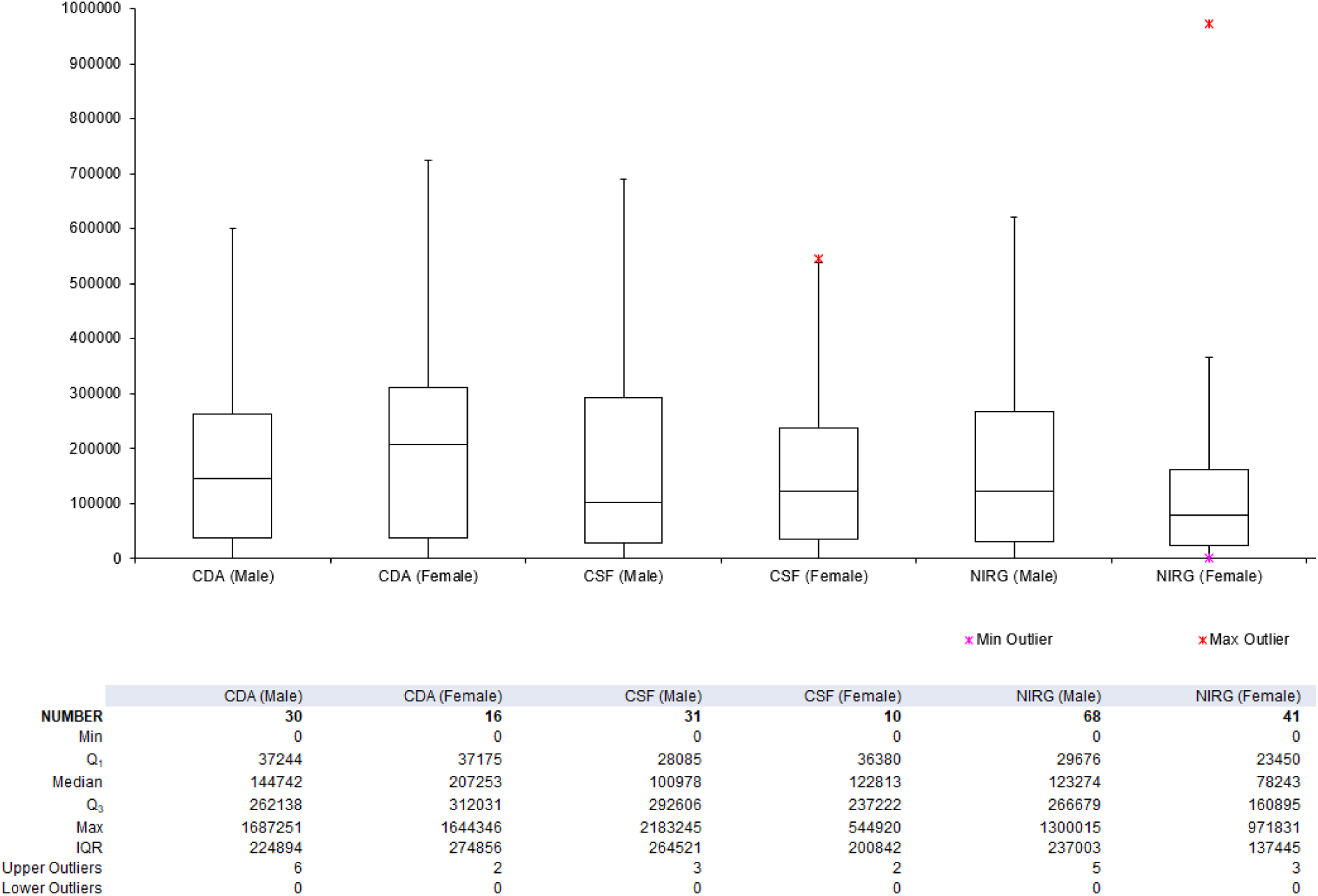
Comparison of average follow-on funding for former NIRG/CDA/CSF holders for three years immediately following completion of the MRC award, by gender

**Figure 5.**
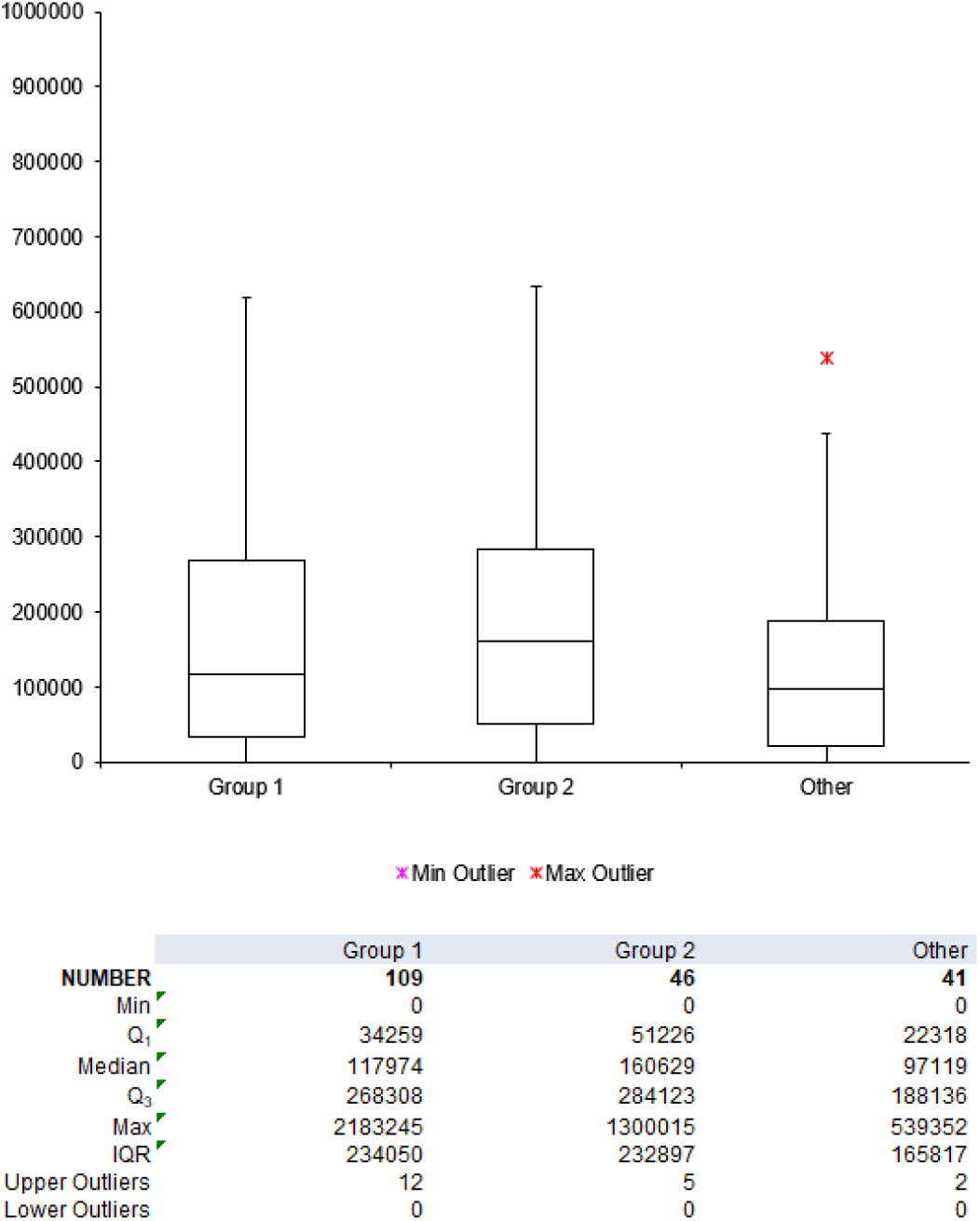
Comparison of average follow-on funding for former NIRG/CDA/CSF holders for three years immediately following completion of the MRC award, by research organisation grouping

### Interviews with ex-NIRG holders confirmed that we had compiled accurate funding details and provide insight into the factors that support career progression

Our interviews with ex-NIRG holders confirmed that we had in most cases accurately captured details of their research income and where there were gaps in this record these were not material (i.e. the information supplied during interview would not have moved the researcher into the next research income category in Figures 1 and 2). All interviewees were actively seeking further research funding. The interviews allowed us to qualitatively explore some of the challenges that NIRG holders faced in securing subsequent grant support. We interviewed 31 researchers (13/34 in our £0-£50,000 per year research income category, 13/21 in our £50,000 to £100,000 per year research income category and 5/41 in the £100,000 to £400,000 per year category).

Factors that had an important impact on progression raised by interviewees included personal circumstances (career breaks, moving institution), support provided by the host institution (bridging funds, start-up packages, core-funded technician/postdocs and students), mentoring and contact with the MRC, as well as some project specific factors (such as interruption of access to infrastructure and recruitment/staffing issues).

- Although the numbers were small^18^, there were examples of career breaks being taken by NIRG holders that subsequently secured high and low research income levels.
- The majority of NIRGs interviewed reported good support from their institution including making available local funding and studentships.
- Finding a mentor and/or supportive network of peers was most widely reported as a factor that had contributed to success (by those with high and low research income) and those researchers unable to secure a helpful mentor as potentially limiting to their career (again by those with high and low research income).
- NIRG holders were positive about contact with the MRC and wanted more opportunities for advice and networking. Those that had been able to take up the opportunity to attend a research board meeting as an observer found this very helpful. Those that had not been approached by the MRC felt that they would have benefited from this contact.

## Conclusions

This study examined in detail one indicator of establishing an independent research programme, specifically the research income secured by early career researchers. There are of course many other aspects that are important such as the quality and productivity of research output, management and mentoring of staff, contribution to the community as a peer reviewer etc. To allow results to be compared easily between cohorts that completed their MRC early career researcher award in different years, and to cope with complex portfolios of follow-on grants with overlapping tenure, we produced an estimate of average research expenditure for the three years immediately following the early career award. This is the first time that a study has been made of the research income secured from all sources for such a cohort of researchers. We were fortunate that almost all researchers in the cohort remained in academic posts and so it was possible to access information about their research income. Our approach to compile details of research income was validated in a small number of interviews with ex-NIRG holders based in the UK. It was acknowledged that details of funding for the 10 per cent of researchers that had moved overseas was likely to be partial and that there was no information about the career for the one per cent that had moved into the private sector.

Although the CDA/CSF/NIRG supported researchers secure similar amounts of grant funding, these awards may offer complementary routes to an independent research career for researchers. Early career researchers applying for NIRGs are prioritised for consideration by the research boards, but NIRGs are typically shorter in duration, have a focus on a single project, and since 2016 only include part of the principal investigator’s salary. Fellowships are highly competitive^19,^ provide one to two years more support than a NIRG plus all the principal investigator’s salary to approach an ambitious programme of work, they therefore require a larger investment from the MRC. The NIRG route requires that researchers combine the MRC award with securing funding from elsewhere from an early point whereas for the most competitive applicants, MRC fellowships provide a little protected time before applying for follow-on funding. Although we found a slightly higher proportion of fellowship holders than NIRG holders secured very substantial levels of research income^20^, the difference was found to be non-significant and there was no clear difference in the proportion that led grants as principal investigator.

We noted that for clinical researchers, fellowships are a more established route to funding than NIRGs. Just eight per cent (nine researchers) of NIRGs supported clinically qualified researchers. There were 41 Clinician Scientist Fellowships in the same sample period. In our small sample of 46 research grants 10 principal investigators were clinically qualified (22 per cent).

We suggest that these data show reasonably good overall progression rates for MRC supported early career researchers, with both MRC NIRGs and MRC fellowships effective in supporting research careers. We found that 50-59 per cent of researchers were expending more than £100,000 per year on their research programme within three years of completing their MRC award. This level of research funding suggests income from a variety of sources, and Universities often cite income in this range as a pre-requisite for being considered for tenured positions.

For PLT researchers the advantage of substantial core funding, a six-year period of support, and the opportunities for progression to a MRC programme leader position, were clear. 94 per cent of ex-PLT researchers were expending more than £100,000 per year on their research programme in the three years after completing their PLT position. This appeared to be the most secure route to an independent research career.

We identified researchers that were of similar age to the CDA/CSF/NIRG holders, without support from these MRC early career funding schemes, that had secured research grants of similar value as a comparator group. We found this group to be highly competitive for follow-on research funding. 89 per cent of these researchers were expending more than £100,000 per year on their research programme in the following three years. We recognise that age is not a good proxy for career stage and suggest that these researchers were likely to have already established their research career or were at an advanced stage of doing so.

There is some evidence of a “bottleneck” to progression with a third of early career researchers from the CDA/CSF/NIRG schemes taking more than three years to secure substantive follow on funding. We only investigated the NIRG cohort further and the researchers interviewed cited a variety of factors that may impact on their progression, including for NIRG holders the increased competition for funding at this next career stage. Across the portfolio the fact that only around a third of early career researchers are still to establish a substantively funded research programme three years after their CDA/CSF/NIRG funding, but almost all are still working toward this, might be considered an indicator that the MRC took an appropriate level of risk in funding these researchers in a highly competitive funding environment. However, for specific research areas in which the MRC wishes to build research capacity, this level of progression would be a barrier to the MRC achieving this goal.

In the three years immediately following completion of an MRC early career award, we found that the MRC is still the most important follow-on funder, but a very wide range of funding sources are used. These results may be helpful in estimating the downstream funding demands that researchers at a similar career stage will place on the research base. We highlight this as initiatives such as the current UK Research and Innovation (UKRI) future leader fellow scheme^21^ are aimed at increasing the population of early career stage researchers. Our results suggest that these researchers may secure approximately three times the value of their early career award in grant support in the three years following their early career award. Additional high-quality demand will require an increase in grant funding beyond the early career stage (e.g. toward research leadership^22^) or success rates will decline and the bottleneck for early career researchers to progress will worsen. Expansion of the research base needs balanced new investment across schemes and stages and cannot be driven through investment in fellowships and NIRGs alone.

Progression of researchers funded via NIRGs and Fellowships was not noticeably influenced by the prominence of the employing research organisation, as measured by MRC research income. The progression rates for researchers based in the MRC’s “top 10”, “top 20” and “all other” research organisations were similar. No statistically significant differences were found between the progression (as measured by average research income) of male and female researchers supported under each scheme, but non-significant differences were noted. NIRG holders that took a career break during or shortly after their MRC award described this as a challenge to maintaining momentum in their research career. However, we have not found evidence that this can explain the small difference in male and female median research income for NIRGs. It was noted that our results also showed male CDA holders had slightly lower median income than female CDA holders, suggesting that the small variations between male and female researchers in our data may be due to chance.

The MRC has, in part as a result of this study and its continuous discussion with grant-holders, implemented several changes to the way it supports early career researchers. In 2019 the MRC launched a £4 million pilot scheme to provide additional support to existing CDA and CSF holders to help them transition to leadership positions^23^. In 2020 the MRC launched a short film^24^ to help explain how the NIRG benefits early career researchers, how to apply and what makes a good application. The MRC has also enhanced opportunities for mentoring^25^, and extended invitations to its in-house fellowship induction meeting to NIRG holders.

The interviews with ex-NIRG holders were illuminating and emphasised that while a useful start, measuring research income took a very narrow view of research career progression. All the ex-NIRG holders were actively pursuing their ambition to establish an independent and substantive research programme, as well as teaching and supervising students. We are interested in expanding this work through further interviews and analysis, to examine other indicators for research progression, and to provide comparisons with non-MRC funded early career researchers.

## Acknowledgements

The authors are all employed in the UKRI-MRC analysis and evaluation team which aims to provide robust evidence for decisions about the support for health research. We would like to thank the interviewees for their time and feedback. We would also like to thank Dr Declan Mulkeen and colleagues in the MRC Training and Careers group for their feedback on earlier drafts of this paper.

